# Global Patterns in Culturable Soil Yeast Diversity

**DOI:** 10.1101/2021.05.19.444851

**Authors:** Himeshi Samarasinghe, Yi Lu, Renad Aljohani, Ahmad Al-Amad, Heather Yoell, Jianping Xu

## Abstract

Yeasts, broadly defined as unicellular fungi, fulfill essential roles in soil ecosystems as decomposers and nutrition sources for fellow soil-dwellers. Broad-scale investigations of soil yeasts pose a methodological challenge as metagenomics are of limited use on this group of fungi. Here we characterize global soil yeast diversity using fungal DNA barcoding on 1473 yeasts cultured from 3826 soil samples obtained from nine countries in six continents. We identify mean annual precipitation and international air travel as two significant predictors of soil yeast community structure and composition worldwide. Anthropogenic influences on soil yeast communities, directly via travel and indirectly via altered rainfall patterns resulting from climate change, are concerning as we found common infectious yeasts frequently distributed in soil in several countries. Our discovery of 41 putative novel species highlights the need to revise the current estimate of ~1500 recognized yeast species. Our findings demonstrate the continued need for culture-based studies to advance our knowledge of environmental yeast diversity.

## 1. Introduction

Soil is host to an incredible amount of microbial life, with each gram containing over 10 billion cells of bacteria, archaea and fungi (Roesch et al., 2007). Despite their relatively low abundance, soil fungi fulfill essential roles in decomposition of organic material, nutrient cycling and soil fertilization (Frac et al., 2018). This is especially true for yeasts, broadly defined as unicellular fungi, whose numbers rarely exceed thousands of cells per gram of soil. Yet, yeasts in soil ecosystems are essential decomposers and nutrient sources for fellow soil-dwelling protists, bacteria, insects, and nematodes (Botha, 2011; Yurkov, 2018). In fact, yeasts may be the predominant soil fungi in cold biospheres such as continental Antarctica (Connell et al., 2008; Vishniac, 1996). Soil is also a primary reservoir for common pathogenic yeasts of the *Candida* and *Cryptococcus* genera that frequently cause superficial and systemic infections in humans (Kurtzman et al., 2011).

It is becoming increasingly apparent that the true extent of global soil yeast diversity is significantly underestimated. While yeast cells were first observed under the microscope in 1680 by Anton Van Leeuwenhoek, their natural habitats were a topic of contention among mycologists who often associated yeasts with fruit trees and fermentation. It was not until the 1950s that soil was established as a true natural habitat of yeasts where they live and reproduce. The culturing media, incubation temperatures and techniques used in pioneering studies were expanded in later projects to isolate soil yeasts of diverse metabolic and functional profiles (reviewed in Yurkov, 2018). Currently, the diverse array of yeasts recovered and characterized from soils across the globe contribute to the ~1500 recognized yeast species on the planet (Kurtzman et al., 2011). Environmental surveys of yeasts routinely uncover novel species, accounting for as much as 30% of yeast populations, highlighting the need to revise current estimates of global yeast diversity (Groenewald et al., 2018; Yurkov et al., 2016b, 2016a).

Lack of adequate environmental sampling, especially in Asia, Africa, South and Central America, limits the discovery of novel yeast species, characterization of soil yeast communities, and prediction of global diversity patterns. Soil yeast populations often differ in structure and composition between locations. With the exception of a few genera that are widespread in soil such as *Cyberlindnera*, *Schwanniomyces*, *Naganishia*, *Goffeauzyma* and *Solicoccozyma* (Botha, 2011), most yeast species have a fragmented distribution with few shared species between sites, even within the same geographical region. One study found that only eight of the 57 species found in soils of Mediterranean xerophyl forests were shared between the three sampling plots in the same locality (Yurkov et al., 2016a). Another study found only a single species to be present in all three sampled temperate forests in Germany (Yurkov et al., 2012). So far, most environmental surveys reported in the literature have been ecologically/ geographically limited, with sampling often focus on a specific ecological niche within a single locality, region, or country (Into et al., 2020; Li et al., 2020; Monteiro Moreira and Martins do Vale, 2020; Tepeeva et al., 2018).

Factors affecting soil yeast diversity have not been fully elucidated but soil moisture, soil pH, carbon content and nitrogen content have been implicated as contributing variables (reviewed in Botha, 2011). In 2006, Vishniac analyzed prominent yeast species in soil along a latitudinal gradient and found that mean annual temperature, mean annual rainfall and electrical conductivity explained ~44% of the variation in yeast species distributions (Vishniac, 2006). As their sampling locations were limited to the Americas and Antarctica, it is unknown whether the same trends persist on a global scale. One potential contributing factor to yeast species and genotype distributions is anthropogenic influences such as international travel. For example, international travel has increased exponentially in the last few decades with direct implications for the global spread of organisms, most notably infectious disease agents and invasive species. The role of global travel in introducing infectious diseases to new areas and facilitating epidemics is well documented (Findlater and Bogoch, 2018), with the current COVID-19 pandemic being a prime example. International travel is likely affecting soil yeast communities by transferring previously geographically isolated species and genotypes across borders, although a link between the two has not been previously investigated.

Metagenomics is widely applied in the study of environmental microbes to investigate taxonomic diversity, characterize functional groups, and elucidate broad scale patterns (Abbasian et al., 2016; Abia et al., 2018; Egidi et al., 2019; Li and Qin, 2005). However, metagenomics and other culture-independent methods cannot be readily applied to the study of yeasts due to the lack of a suitable yeast-specific barcoding gene (Xu, 2016). Yeasts are phylogenetically diverse and occur among filamentous fungi in two major phyla, Ascomycota and Basidiomycota, within the fungal kingdom. In a 2014 study that has not been surpassed in scale before or since, Tedersoo and colleagues used high throughput sequencing of fungal barcoding DNA to assess global soil fungal diversity and identify predictors of global diversity patterns (Tedersoo et al., 2014). Due to the lack of a sequence-based signature, yeasts were not singled out as a group of interest and thus limited information was presented on soil yeast diversity of the 39 countries included in the study. They identified mean annual precipitation and distance from equator as the two strongest overall predictors of soil fungal diversity on a global scale. It is not clear if and to what extent the same predictors apply to yeast diversity in soil.

Using a global collection of 3826 soil samples, here we assessed the culturable soil yeast diversity in nine countries representing all continents except Antarctica. We found soil yeast populations of each country to be unique in structure and composition as 73% of the discovered species were not shared between countries. Mean annual precipitation was the most significant predictor of culturable soil yeast diversity on a global scale. We found air traffic volume to be significantly correlated with the number of shared species between countries, implying a link between international travel and transfer of yeast species across borders. Our study overcomes the geographical constraints of many previous studies by identifying soil yeast diversity patterns on a global scale. We also demonstrate that culture-dependent methods provide a more comprehensive framework than metagenomics for studying phylogenetically diverse, but morphologically targeted groups of organisms such as yeasts.

## 2. Materials and Methods

### 2.1 Soil collection

We collected soil from 53 locations in nine countries encompassing all continents except Antarctica (Figure 1 and Supplementary Table 1). At each location, bags (3cm x 7cm sterile, resealable, polyethylene bags) of topsoil, within 1-3 inches from the surface, were collected following sterile protocols and transported to our lab at McMaster University, Canada. Each bag contained soil from 10 sites in the same area but located at least 2m from each other. Different bags represented soils from at least 100m from each other. The soil in each bag was segregated into ~1g aliquots and stored at 4°C. In total, this study included 3826 soil samples originating from the following countries: Cameroon (493), Canada (300), China (340), Costa Rica (388), France (327), Iceland (316), New Zealand (610), Peru (490) and Saudi Arabia (562).

**Figure 1:**
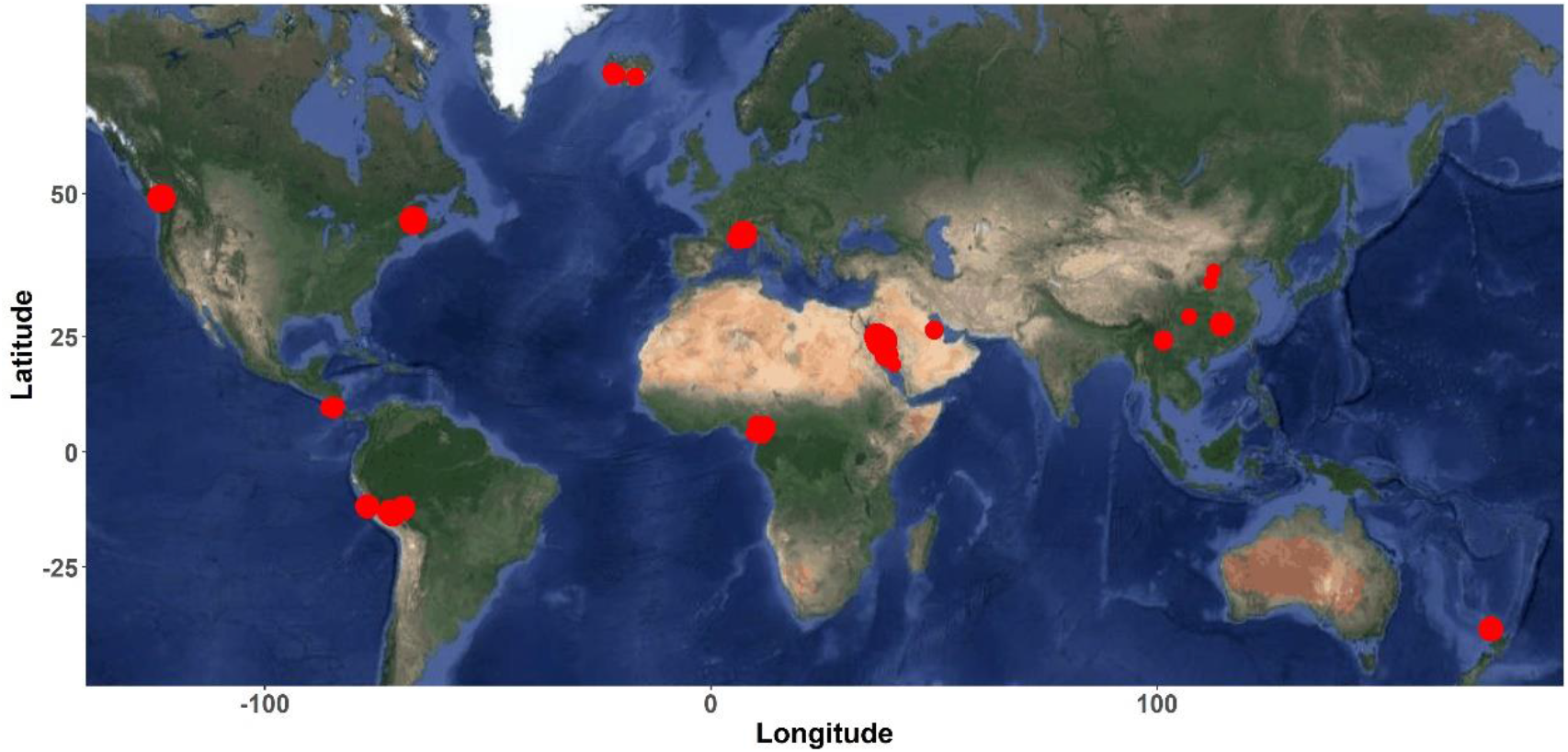
Soil sampling locations. Soil was collected from 53 locations, indicated by the red circles, in nine countries. The size of the circle corresponds to the number of samples obtained from that location.

### 2.2 Yeast isolation from soil samples

Yeasts were isolated at a temperature deemed to be optimal based on the country of origin’s mean annual temperature. For each soil sample, approximately 0.1g was added into 5ml of YEPD broth (Yeast Extract-Peptone-Dextrose) in 13ml culture tubes and incubated in a roller drum for 24 hours. The broth contained the antibiotic chloramphenicol (50mg/L) and the selectively toxic fungicide benomyl (5mg/L) to inhibit bacterial and mold growth, respectively. We extended this incubation step to 72 hours for soil samples from Iceland due to slower yeast growth at 14°C. We then plated 100ul of the broth onto solid YEPD containing chloramphenicol and benomyl and incubated at the same temperature for an additional 2-5 days until microbial growth was visible. For each plate that contained morphologically yeast-like colonies, we randomly selected a representative colony and streaked it onto fresh YEPD plates to obtain single colonies. If more than one morphology was present, one representative colony of each type was separately streaked for single colonies. After 2-3 days’ incubation, we randomly picked one single colony per yeast isolate and suspended in 50ul nuclease-free water to be used in <u>P</u>olymerase <u>C</u>hain <u>R</u>eaction (PCR).

### 2.3 Yeast identification via ITS sequencing

We identified the yeasts by sequencing their fungal barcoding gene, the ribosomal <u>I</u>nternal <u>T</u>ranscribed <u>S</u>pacer (ITS) regions. We performed colony PCR using primers ITS1 (5’ TCCGTAGGTGAACCTGCGG 3’) and ITS4 (5’ TCCTCCGCTTATTGATATGC 3’) to amplify the ITS region. The PCR cocktail consisted of 10ul Promega GoTaq Green Mastermix, 5ul nuclease-free water, 2ul each of the primers (2uM) and 1ul cell suspension. The thermocycling conditions were an initial denaturation step at 95°C for 10 minutes followed by 35 cycles of (i) 95°C for 30 seconds, (ii) 55°C for 30 seconds and iii) 72°C for 1 minute. We ran 4ul of the PCR products on a 1.5% agarose gel to check for successful amplification. The remaining PCR products were Sanger sequenced with the ITS1 primer at Eurofins Genomics in Louisville, Kentucky (https://eurofinsgenomics.com/en/home/). We trimmed the low-quality ends of the ABI chromatograms generated from Sanger sequencing and batch converted to FASTA format with DNA Baser’s ABI to FASTA converter software (https://www.dnabaser.com/download/Abi-to-Fasta-converter/abi-to-fasta-converter.html). We used BLAST+ applications on the command line to query the multi-FASTA file against the NCBI nucleotide database to detect sequence similarity to existing ITS sequences. The BLAST searches were run remotely (-remote flag) to avoid downloading the entire database onto our servers. The output was compiled into a CSV (Comma Separated Values) file containing the top 10 matches for each query sequence. We inspected the CSV file manually to check for quality and for any inconsistencies in species identity within the top ten matches. The putatively identified species through GenBank searches were further confirmed by manually comparing to sequences in the curated UNITE database (https://unite.ut.ee/). We assigned species identities to our ITS sequences at a sequence similarity threshold of 98.41% to existing sequences in databases. This threshold was previously determined to be optimum to distinguish yeast species at the ITS locus based on an analysis of 9000 fungal sequences (Vu et al., 2016). Sequences with no matches surpassing this threshold were considered putative novel species.

### 2.4 Statistical analyses of population diversity

All statistical analyses were conducted in RStudio v.4.0.2 using a combination of base functions and packages including ggmap (Kahle and Wickham, 2013), ggplot2 (Wickham, 2016), and tidyverse (Wickham et al., 2019). We quantified the diversity of yeast populations at our sampling sites by calculating the Shannon diversity index using the package Vegan v.2.5-7 (Oksanen et al., 2020). We conducted rarefaction analyses using the iNEXT package (Hsieh et al., 2016) to determine if sufficient soil sampling was performed in each of the nine countries to accurately estimate their culturable soil yeast diversity.

### 2.5 Relationship between yeast diversity and climate and geographic factors

Within each country, soil collection sites differed in climatic and environmental conditions, with the exception of New Zealand where sampling was limited to the metropolitan region of Auckland. We assigned the 53 sampling sites to 47 distinct locations based on their geographical coordinates. Using geographical coordinates, we calculated mean annual precipitation and mean annual temperature by averaging monthly data over a 16-year period from 1991-2016, available on Climate Change Knowledge Portal (https://climateknowledgeportal.worldbank.org/). We calculated the elevation and distance from the sampling sites to the equator using Google Maps. We calculated the Shannon diversity index of the yeast populations found at the 47 distinct locations. We constructed mixed models using the package lme4 v.1.1-26 (Bates et al., 2015) where precipitation, temperature, elevation, and distance to equator were set as fixed effects, country was fitted as a random effect and Shannon diversity Index was fitted as the dependent variable.

### 2.6 Air traffic data

We extracted data on the number of flights occurring between each country-country pair over a 5-year period from 2011-2016 from the Global Transnational Mobility Dataset (Recchi et al., 2019). This dataset is compiled based on a combination of tourism data and distance-adjusted air-traffic data. Next, we calculated the number of yeast species shared between each country-country combination. To assess the correlation between the number of shared species and the volume of air traffic between countries, a linear model was fitted between the two variables.

### 2.7 Comparison to metagenomics study

We compared our findings to a previous study that used culture-independent methods to investigate global diversity of soil fungi. In 2014, Tedersoo and colleagues extracted DNA directly from soil samples of 39 countries and performed high throughput sequencing of the ITS2 region using primers ITS3 and ITS4 (Tedersoo et al., 2014), covering a portion of the DNA barcoding fragment we sequenced here. Four countries overlapped between the two studies, namely, Cameroon, Canada, China and New Zealand. For each of the four countries, we performed BLAST searches using BLAST+ applications on the command line to identify ITS sequences that appeared in both studies. First, we created custom databases containing our ITS sequences for each country. Next, the ITS sequences for the countries of interest were extracted from Supplementary data table 1 of Tedersoo et al.’s paper and converted to FASTA format. The multi-FASTA file for each country was then queried against its respective custom database using the blastn option. The output was compiled into a CSV file containing the top 5 matches for each query sequence. This CSV file was perused manually to identify significant sequence similarity between query and match sequences.

**Table 1:** Summary statistics of yeast isolation from global soil samples

| Country | Number of soil samples | Number of yeast isolates | Number of known species | Number of novel species | Number of country-specific species | Shannon diversity index |
| --- | --- | --- | --- | --- | --- | --- |
| Cameroon | 493 | 110 | 10 | 9 | 12 | 2.17 |
| Canada | 300 | 261 | 34 | 12 | 25 | 3.06 |
| China | 340 | 230 | 23 | 5 | 15 | 2.54 |
| Costa Rica | 388 | 95 | 20 | 2 | 9 | 2.21 |
| France | 327 | 175 | 12 | 2 | 3 | 1.26 |
| Iceland | 316 | 211 | 11 | 0 | 4 | 1.25 |
| New Zealand | 610 | 155 | 14 | 4 | 5 | 2.05 |
| Peru | 490 | 139 | 30 | 9 | 20 | 3.27 |
| Saudi Arabia | 562 | 97 | 8 | 1 | 5 | 0.91 |
| **Total** | **3826** | **1473** | **90** | **44** | **98** |  |

## Results

### 3.1 Yeast isolation and species identification

We isolated a total of 1473 yeasts from 3826 soil samples (Table 1). The isolation rate varied among the countries, ranging from 17% in Saudi Arabia to 87% in Canada. The yeast isolation and species distribution data from Cameroon soils have been reported in a previous study (Aljohani et al. 2018). Overall, we observed a slightly negative correlation between the number of soil samples and the yeast isolation rate (p < 0.05), as countries with more soil samples did not necessarily yield more yeast isolates.

We successfully assigned species identity to 1367 isolates using the 98.41% sequence identity cut-off to homologous ITS sequences in NCBI and UNITE databases. These strains were categorized into 90 species belonging to 37 genera, with *Candida* being the most species-rich genus (n= 19 species). With 60 ascomycetes and 30 basidiomycetes, both major yeast-containing phyla within the fungal kingdom were broadly represented. The 90 species belonged to six Classes, ten Orders and 18 Families. However, two genera, *Nadsonia* and *Holtermanniella,* do not currently have defined Family associations (*incertae sedis*). The remaining 106 yeast strains can be grouped into 44 unique clusters at 98.41% nucleotide similarity. Since no existing sequences with >98.41% ITS sequence identity were found in the databases, these 44 clusters represent potentially novel yeast species. Genbank accession numbers to the ITS sequences of our 1473 isolates are MG817572 to MG817630 and MW894661 to MW896112 (Supplementary dataset 1).

Our rarefaction analyses suggested that sufficient soil sampling was conducted in each country to accurately estimate the true diversity of culturable soil yeasts. Projections for Shannon diversity index beyond the number of soil samples included in the study revealed that the diversity of our yeast populations approached saturation asymptote (Supplementary Figure 1). Additional sampling in these locations was not likely to have revealed higher yeast species diversity.

### 3.2 Diversity and abundance of culturable soil yeast populations

The abundance and diversity of soil yeast populations varied significantly between countries. Saudi Arabia ranked lowest among the nine countries in the number of unique species, where 562 soil samples yielded 97 yeast isolates belonging to 9 species, 1 of which was novel. On the other hand, we obtained 261 yeast isolates from 300 Canadian soil samples, encompassing 46 species, 12 of which were novel. The number of yeast isolates and of distinct species found in the seven remaining countries ranged from 95-230 and 11-39 respectively (Table 1). The Shannon diversity index of the soil yeast populations ranged from 0.91 (Saudi Arabia) to 3.27 (Peru). Less diverse populations tended to be dominated by a single yeast species, most notably in France, Iceland, and Saudi Arabia where *Candida subhashii*, *Goffeauzyma gastrica* and *Cryptococcus deneoformans* predominated the soil yeasts respectively (Figure 2).

**Figure 2:**
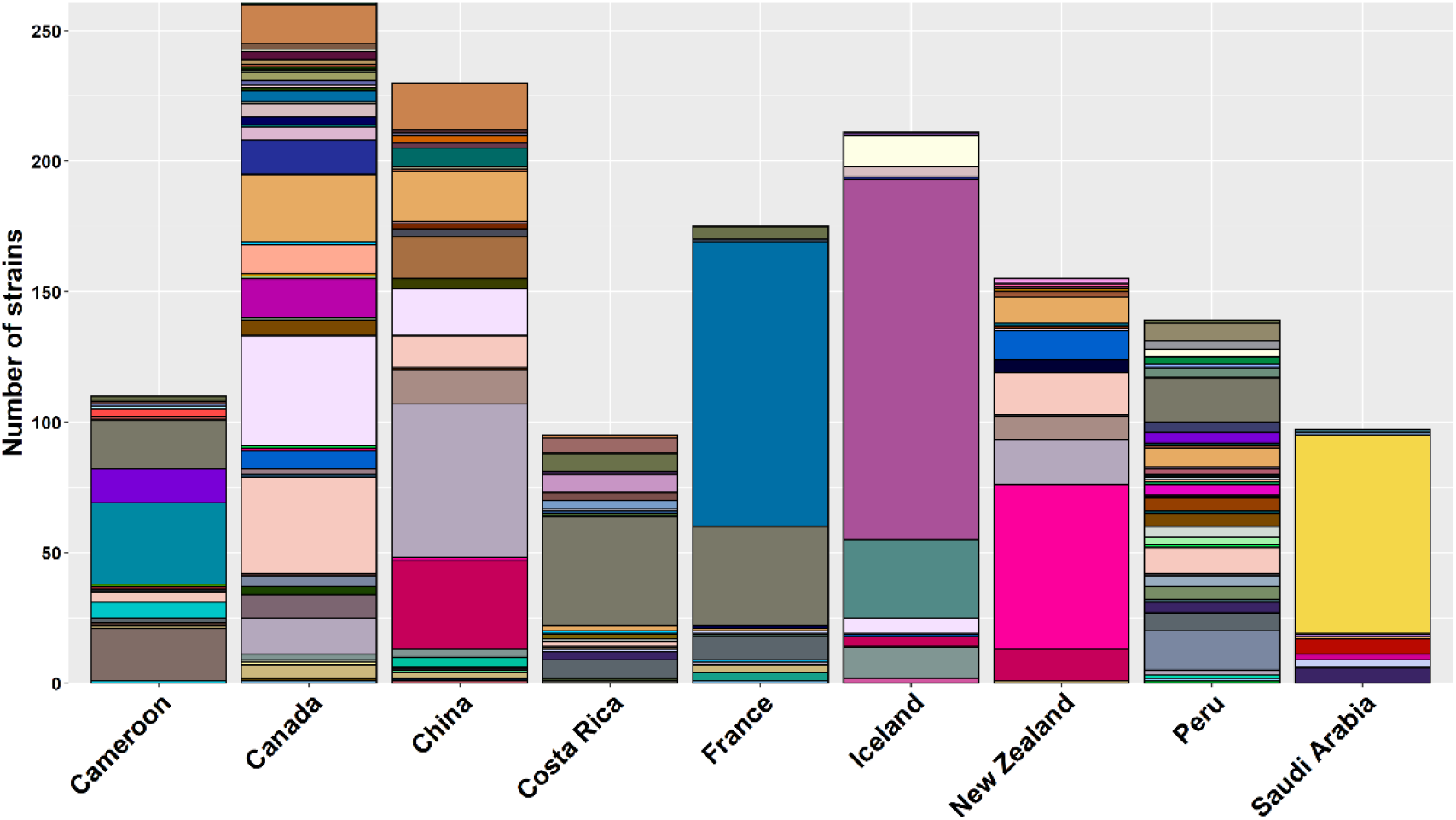
Culturable soil yeast populations by country. The X-axis represents the country. Each country is represented by a stacked bar plot. Each colour represents a unique species and the height of the colored sections indicate the abundance of that species.

#### 3.2.1 Cameroon

As shown in the study by Aljohani et al. (2018) for the Cameroonian soil samples, all but one of the 19 yeast species recovered from Cameroonian soil were ascomycetes. The population was dominated by four species: *Cyberlindnera subsufficiens* (28%), *Torulaspora globosa* (18%), *Candida tropicalis* (17%) and *Cyberlindnera saturnus* (12%). Overall, Cameroonian soil contained the second highest number of novel species (n=9) after Canada (n= 12).

#### 3.2.2 Canada

The culturable yeast population in Canadian soil was dominated by ascomycetes (37 species): the remaining 9 species were basidiomycetes. *Nadsonia starkeyi-henricii,* a little-known yeast that prefers relatively mild temperatures below 25°C, was the most abundant (16%) followed by the pathogenic *Papiliotrema laurentii* (14%). *Debaryomyces hansenii* (10%), *Barnettozyma californica* (6%) and a novel species (6%) were also present in significant amounts. Canadian soil contained 25 yeast species not found in soil samples of the other eight countries, including cold-adapted yeast *Cystofilobasidium ferigula*, industrial lactose fermenter *Kluyveromyces lactis*, and close relative of Baker’s yeast, *Saccharomyces paradoxus.* Canada also had the highest number of novel species (n= 12) of the 9 sampled countries.

#### 3.2.3 China

The Chinese culturable soil yeast population consisted of similar numbers of ascomycetes and basidiomycetes (15 and 13 respectively). This population was dominated by strains of *Solicoccozyma aeria* and *Solicoccozyma terrea* that together accounted for 40% of the population. Other yeasts with significant prevalence included *Debaryomyces hansenii* (8%), *Barnettozyma californica* (8%), *Nadsonia starkeyi-henricii* (8%) and a novel species (7%). 15 yeast species were only found in the Chinese soil which also yielded five novel species.

#### 3.2.4 Costa Rica

*Candida tropicalis* was the dominant species in the culturable soil yeast population of Costa Rica with a prevalence of 44%. The frequencies of the remaining 21 species ranged from 1%-7%. Ascomycetous species outnumbered basidiomycetes at 18 to four. Costa Rica was notable for being the only sampled country to contain strains of the common pathogenic yeasts *Candida albicans* (6%) and *Candida orthopsilosis* (3%). Strains of pathogenic *Candida parapsilosis* were also present in Costa Rican soil (3%).

#### 3.2.5 France

The majority of species in the French culturable soil yeast population were ascomycetes (n= 11) while three species were basidiomycetes. *Candida subhashii*, a pathogenic *Candida* species first identified in 2009 (Adam et al., 2009), was the dominant yeast in this population with an abundance of 62%. *C. subhashii* was previously determined to have strong antagonistic activity against filamentous fungi and has potential as a biocontrol agent against plant pathogenic fungi (Hilber-Bodmer et al., 2017). The widespread pathogen *Candida tropicalis* was the second most abundant species (22%), followed by *Saccharomyces cerevisiae* (5%). One strain of *Candida parapsilosis* was also detected in French soils.

#### 3.2.6 Iceland

Of the 11 species isolated from Iceland soil, six were basidiomycetes and five were ascomycetes. With an abundance of 65%, *Goffeauzyma gastrica* was the dominant species in the Iceland culturable soil yeast population. *G. gastrica* is a cold-tolerant yeast commonly isolated from environmental sources in Antarctica and is known for its production of antifreeze proteins (Białkowska et al., 2017; Ogaki et al., 2020; Villarreal et al., 2018). *Goffeauzyma gilvescens,* another cold-tolerant yeast commonly found in Antarctica, was the second most abundant (14%), followed by *Candida sake* (6%) and *Solicoccozyma terricola* (6%). Iceland was the only sampled country to not yield any novel yeast species.

#### 3.2.7 New Zealand

Basidiomycete species (n= 10) were slightly more prevalent than ascomycete species (n=8) in the New Zealand culturable soil yeast population. *Solicoccozyma phenolica* was the most abundant species with a prevalence of 41%, followed by *Solicoccozyma aeria* (11%), *Papiliotrema laurentii* (10%) and *Solicoccozyma terrea* (8%). We isolated several species with industrial potential from New Zealand soil including *Papiliotrema terrestris*, shown to produce β-galactosidase that was safe for use in food production(Ke et al., 2018), and *Citeromyces matritensis,* an osmotolerant, ethanol-producing yeast shown to be capable of ethanol production from salted algae (Okai et al., 2017).

#### 3.2.8 Peru

Peru’s culturable soil yeast population, consisting of 39 species, ranked the highest among sampled countries in Shannon diversity index. This population was unique in structure and composition as it contained 20 species not found in any other sampled country, including often misidentified pathogen and crude palm oil assimilator *Candida palmioleophila* (Jensen and Arendrup, 2011; NAKASE et al., 1988), rare pathogen *Filobasidium magnum* (Aboutalebian et al., 2020), and halotolerant yeast used in azo dye decolorization *Pichia occidentalis* (Wang et al., 2020). This population contained significantly more ascomycete species (n=29) than basidiomycete species (n= 10). Peruvian population was notable for its relative evenness with no single species exceeding 12% in abundance. *Candida tropicalis* was the most prevalent (12%), followed by *Schwanniomyces occidentalis* (11%) and *Papiliotrema laurentii* (7%).

#### 3.2.9 Saudi Arabia

Saudi Arabian culturable soil yeast population was the least diverse of all sampled countries according to the Shannon diversity index. This population was notable for the overwhelming prevalence of the human pathogenic yeast, *Cryptococcus deneoformans* (78%), the causative agent of fatal fungal meningoencephalitis. The genotypes of C. deneoformans strains from Saudi Arabia have been reported in an earlier study (Samarasinghe et al., 2019). This study was the first to report the environmental presence of *C. deneoformans* in a desert climate. However, overall, the Saudi Arabian soil yeast population consisted of six basidiomycete species and three ascomycete species. One of the species was novel.

### 3.3 Pathogenic yeast species

According to the information presented in the latest edition of The Yeasts: A Taxonomic Study (Kurtzman et al., 2011), the following 12 species were the most common yeast pathogens of humans worldwide: *Candida albicans*, *Candida dubliniensis*, *Candida glabrata*, *Candida guilliermondii*, *Candida krusei*, *Candida lusitaniae*, *Candida parapsilosis*, *Candida orthopsilosis*, *Candida metapsilosis*, *Candida tropicalis*, *Cryptococcus neoformans* and *Cryptococcus deneoformans*. We found 220 strains belonging to eight of these species, accounting for 15% of all yeast isolates found in our samples (Figure 3). *C. tropicalis* was both the most abundant and most widespread with 117 isolates originating from Cameroon, Canada, Costa Rica, France and Peru. The 76 *C. deneoformans* isolates were exclusively found in Saudi Arabian soils. Additionally, seven *C. krusei* isolates were found in Costa Rica, six *C. albicans* isolates were found in Costa Rica, five *C. parapsilosis* isolates were found in Costa Rica, France and Saudi arabia, four *C. lusitaniae* isolates were found in Canada and France, four *C. orthopsilosis* isolates were found in Cameroon and Costa Rica and a single *C. glabrata* isolate was found in Costa Rica. Common pathogenic yeasts were not isolated from our natural soils of China, Iceland and New Zealand.

**Figure 3:**
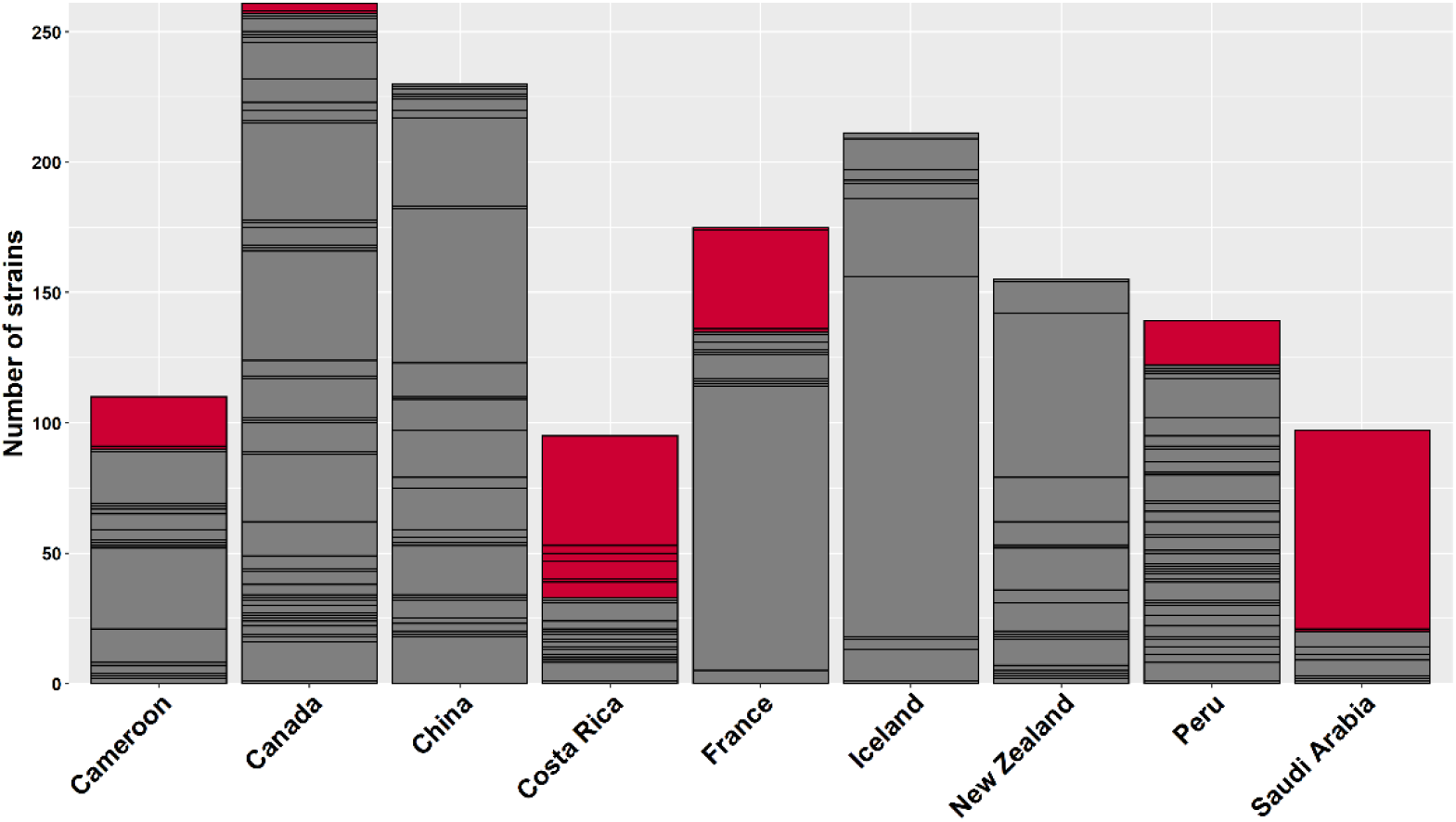
Pathogenic yeast species and their abundance highlighted in red among countries. In these stacked bar plots, the pathogenic species are highlighted in red. The height of the red sections indicates their abundance. Soils from China, Iceland and New Zealand did not yield any pathogenic species.

### 3.4 Novel species

Our yeast population included 44 potentially novel species from eight sampled countries: Our soil samples from Iceland did not yield any novel yeast species. We determined the most closely related genera for 41 species by running BLAST searches in Unite database (remaining three species’ ITS sequences were too short for analysis). Our 41 novel species can be categorized into 12 genera (9 ascomycetes, 3 basidiomycetes) with *Wickerhamomyces* containing ten novel species, and *Candida* containing eight. For each genus, we constructed maximum likelihood (ML) trees using RaxML with 1000 bootstraps (Stamatakis, 2014) to determine the taxonomic placement of novel species with respect to all known species of that genus. Our ML trees confirmed the separation of newly discovered species from known species: for an example, the ML tree of *Wickherhamomyces* species places each novel species at its own distinct node (Figure 4). We observed some geographical clustering where the Cameroonian and Canadian novel species formed their own clusters: the two novel species of Costa Rica and Peru clustered together. The ML trees of the remaining 11 genera can be found in Supplementary Dataset 1.

**Figure 4:**
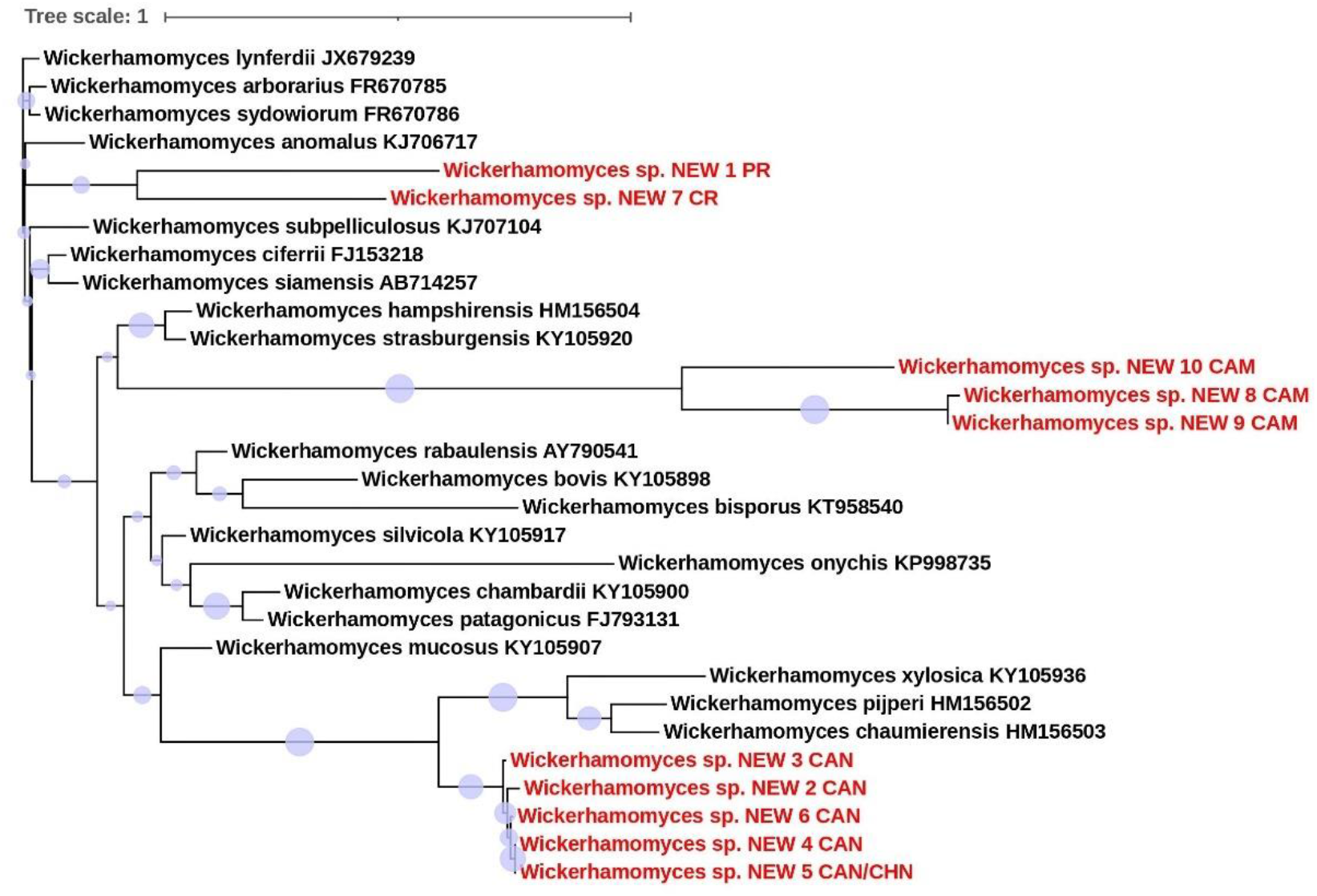
The maximum likelihood tree of *Wicherhamomyces* species. The placement of novel species with reference to known *Wickerhamomyces* species is shown. The novel species’ country of origin is shown in the OTU labels where CAM = Cameroon, CAN = Canada and CHN = China. The tree was constructed using RaxML with 1000 bootstraps.

### 3.5 Predictors of global culturable soil yeast diversity

Our 47 distinct sampling locations covered a wide range of global climatic conditions (Table 2) with mean annual precipitation ranging from 0mm (Lima, Peru where there is virtually no rainfall) to 2965mm (Monteverde, Costa Rica), while mean annual temperature ranged from −1.4°C (Svartifoss, Iceland) to 29.6°C (Alqunfudah, Saudi Arabia). Elevation ranged from < 2m (some sites in Svartifoss, Iceland and Auckland, New Zealand) to 4922km above sea level (Rainbow mountain, Peru). Mbandoumou in Cameroon was the closest to the equator (418.5km from equator) while Dimmuborgir in Iceland was the farthest (7280.5km from equator). Four locations, two in Saudi Arabia and two in Cameroon, were removed from further analysis as they did not yield any yeast isolates. The remaining 43 locations varied significantly in culturable yeast diversity as quantified by Shannon diversity index from 0 (only one species was found) to 2.77 (Fredericton, Canada). According to our mixed model, we found mean annual precipitation to be significantly correlated with the Shannon diversity index (p = 0.012, Figure 5). We found no significant correlation between the remaining predictors and Shannon diversity index (Supplementary Dataset 3).

**Figure 5:**
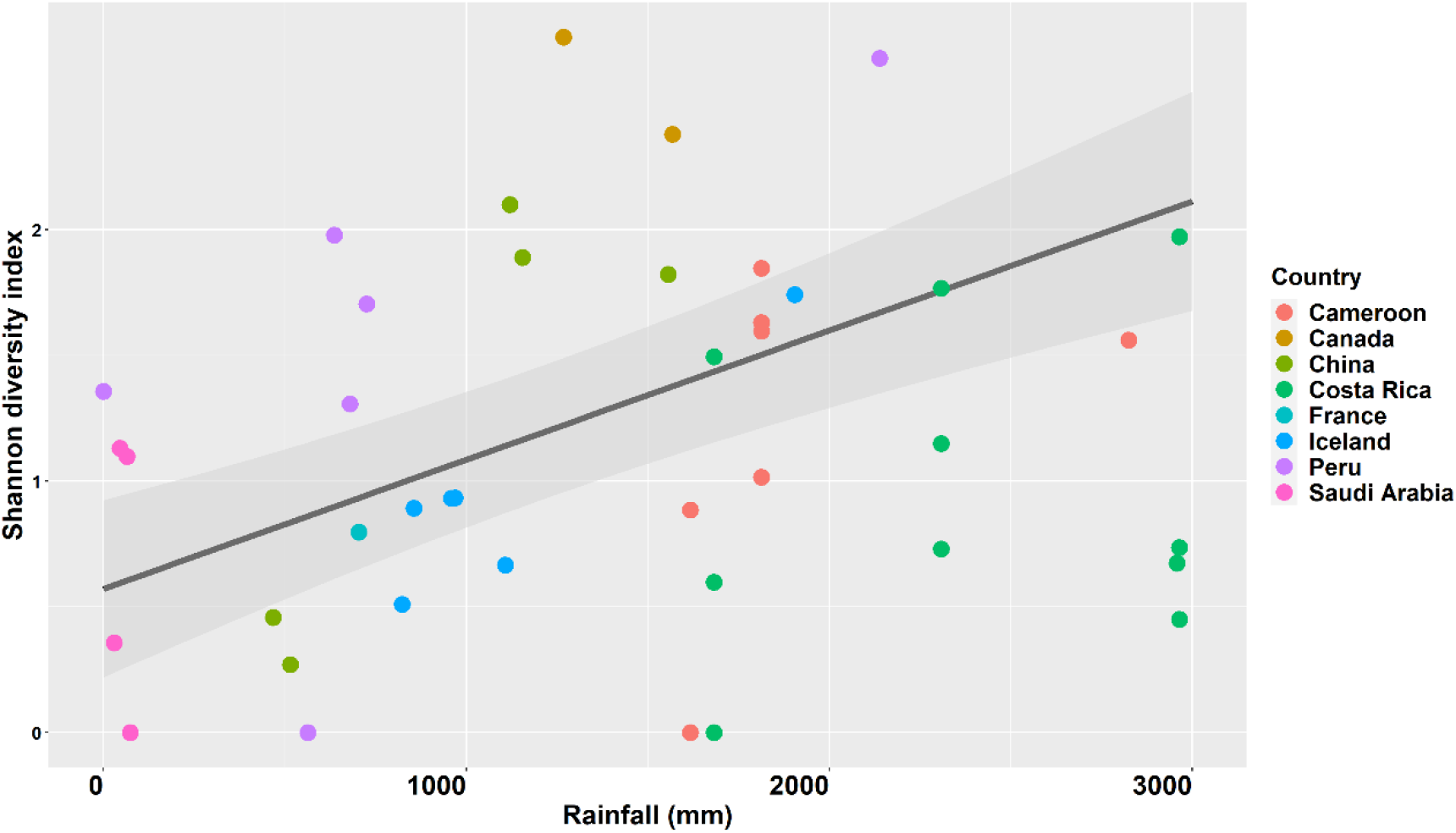
Mean annual rainfall is significantly correlated with soil yeast diversity. Here, the Shannon diversity index of our sampling sites is plotted against mean annual precipitation. Sampling sites are colored by country. The line plots the model predictions with associated uncertainty shaded in grey.

**Table 2:**
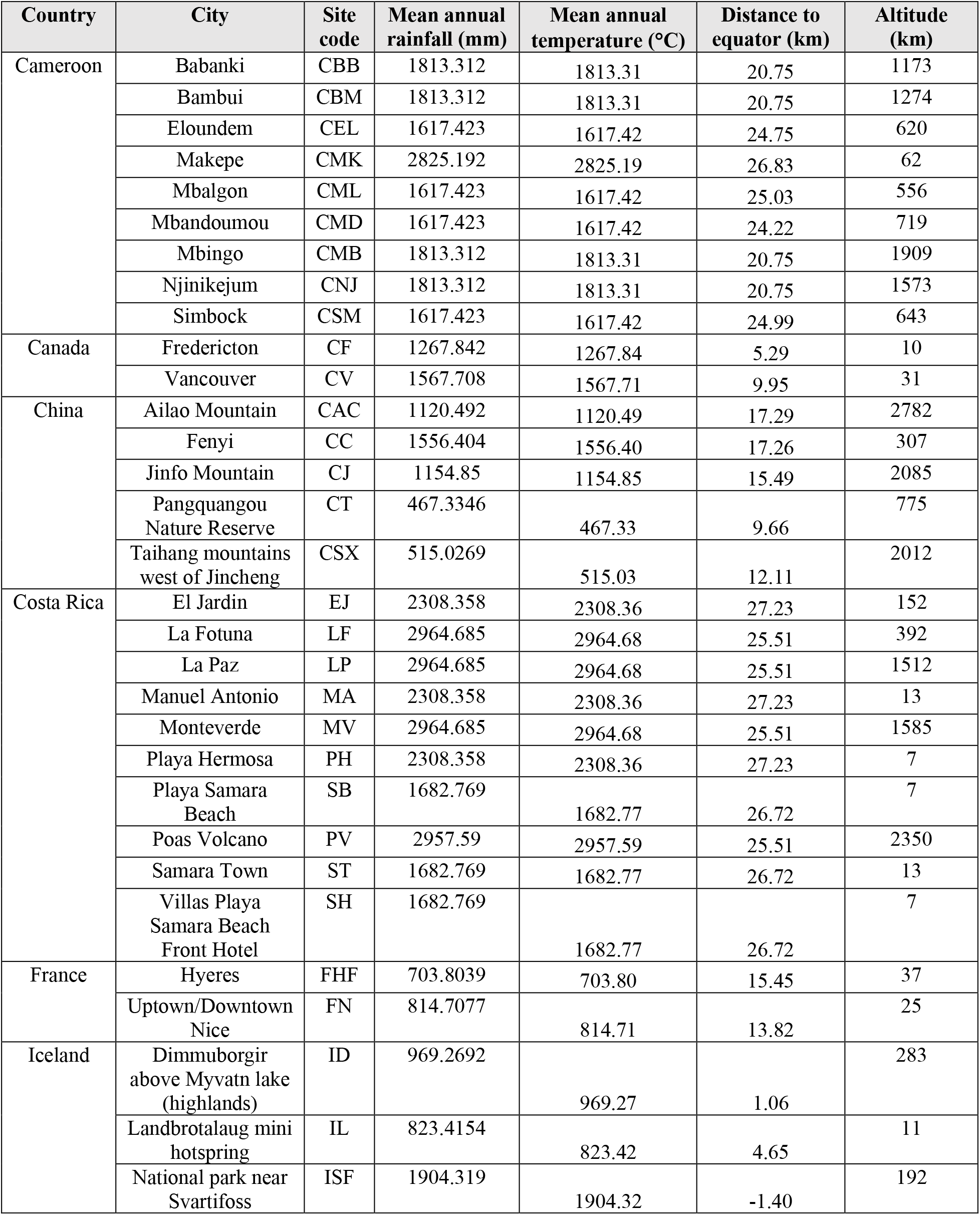
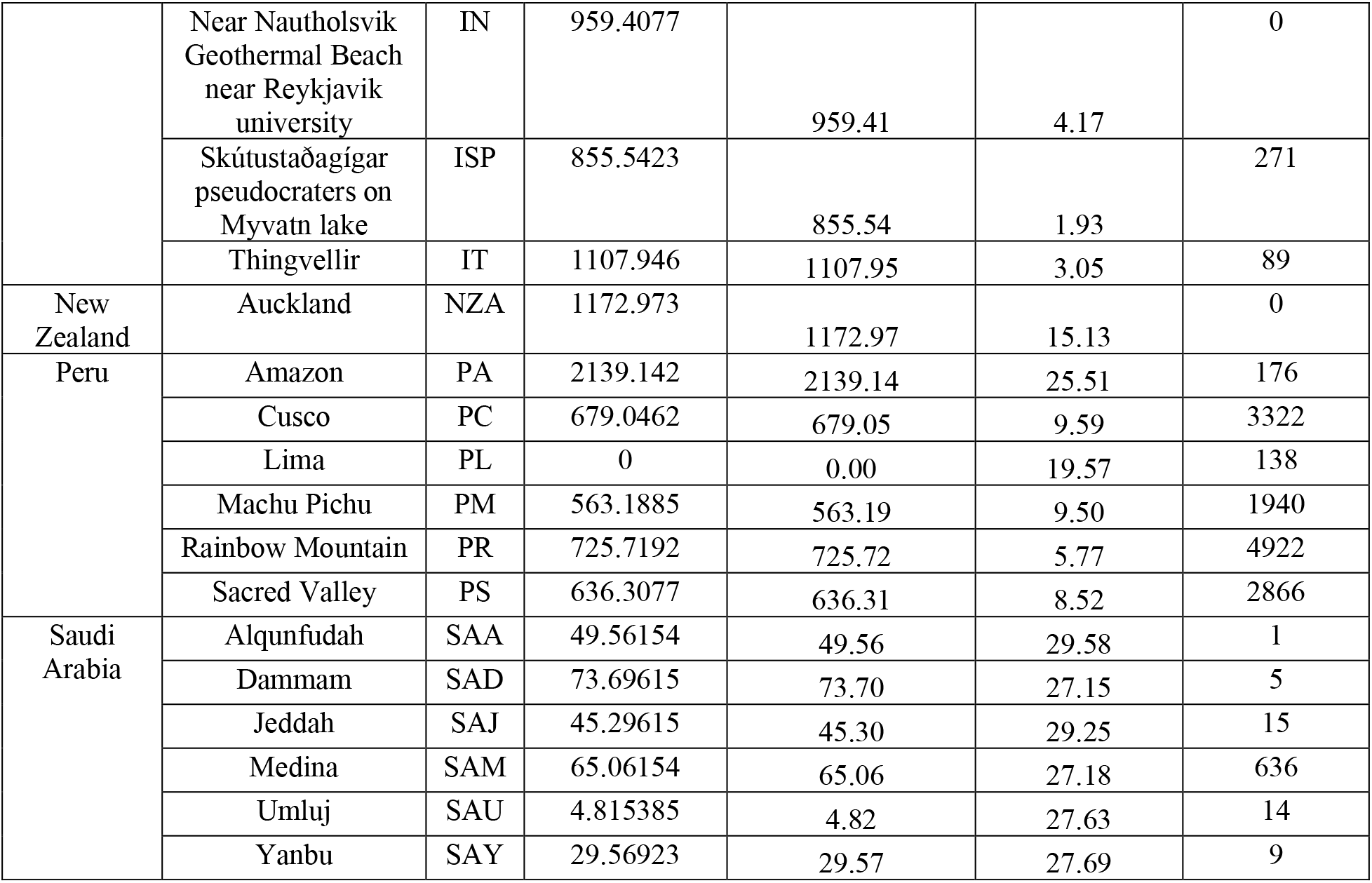
Environmental and geographic characteristics of sampling sites. Mean annual rainfall, mean annual temperature, distance to equator and elevation of sampling sites are summarized here.

### 3.6 Air traffic volume as a predictor of shared species between countries

36 yeast species were found in more than one country. The following five country pairs had no soil yeast species in common: Iceland-Cameroon, Iceland-Costa Rica, Iceland-France, Iceland-Saudi Arabia, and Cameroon-Saudi Arabia. The number of shared species between the remaining 31 pairs ranged from one to 11 (Figure 6b). Air traffic volume data extracted from the Global Transnational Mobility Dataset showed that 25 700 496 trips were made between China and France between 2011-2016. During the same period, only 81 trips were made between Iceland and Cameroon (Figure 6a, Supplementary Table 3). We performed a linear regression analysis between air traffic volume, geographic distance, and the number of shared species between countries. While we found no significant correlation between geographic distance and sharedness, air traffic volume was significantly correlated with the number of shared species between countries (p = 0.003, Figure 7).

**Figure 6:**
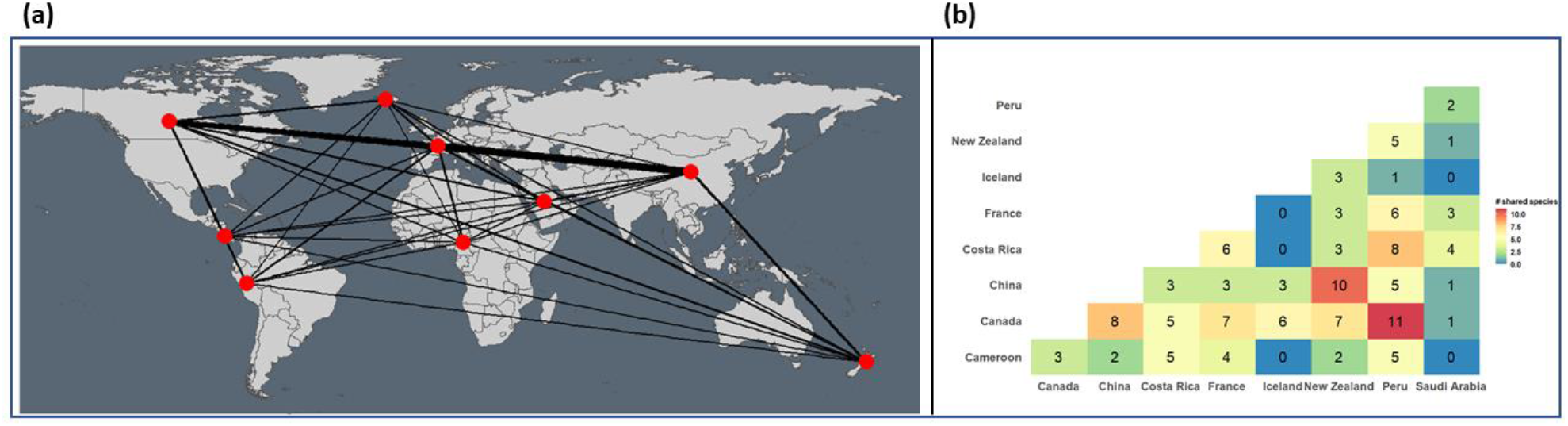
Air traffic volume between countries is correlated with number of shared species. (a) Volume of air traffic between the nine countries from 2011-2016. Thickness of the line indicates volume. (b) Heat map showing the number of shared species between country pairs.

**Figure 7:**
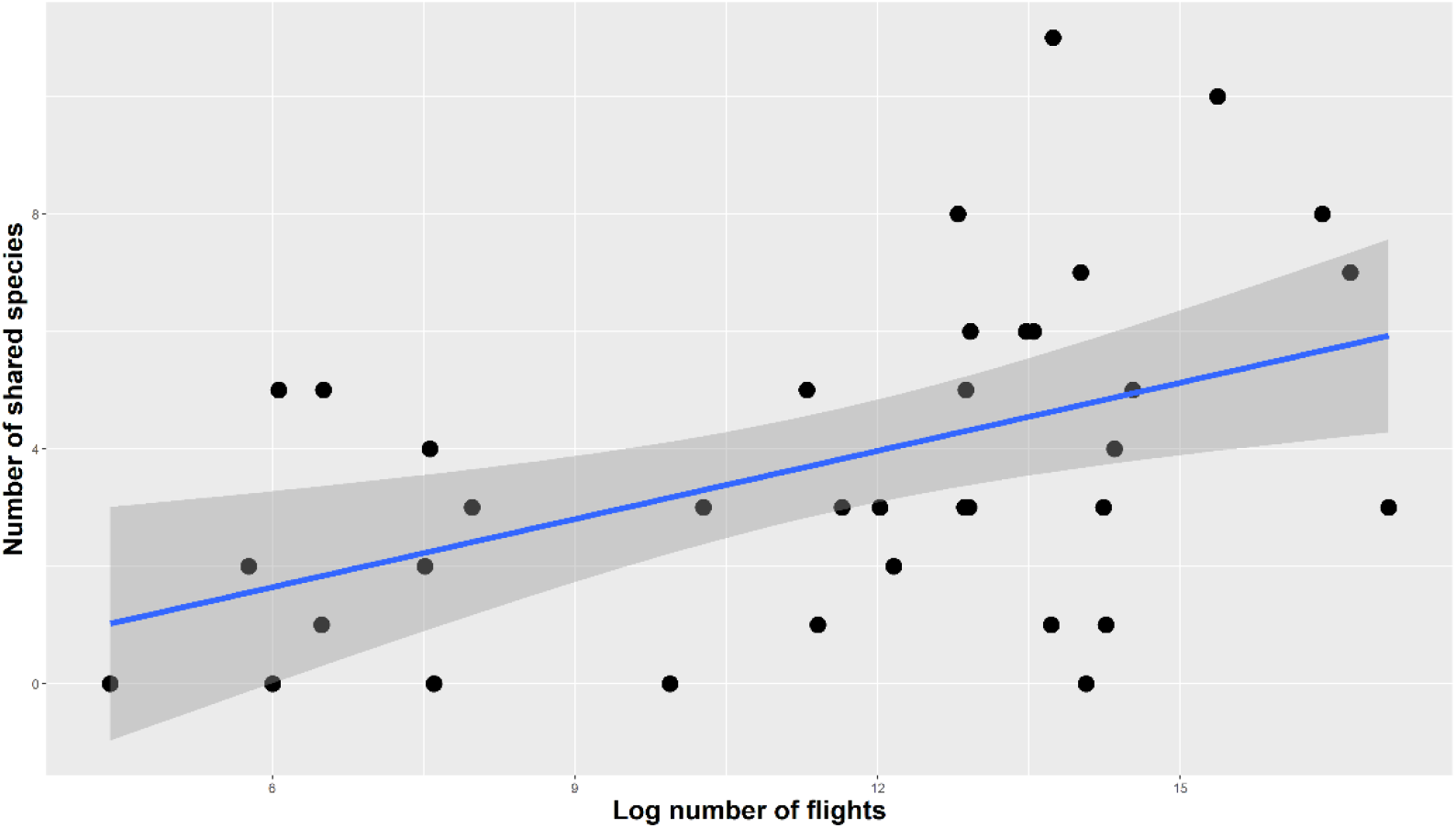
Number of shared species between countries is significantly correlated with traffic volume between them. We plotted the log number of trips made between country pairs between 2011-2016 against the number of yeast species shared between them.

**Figure 8:**
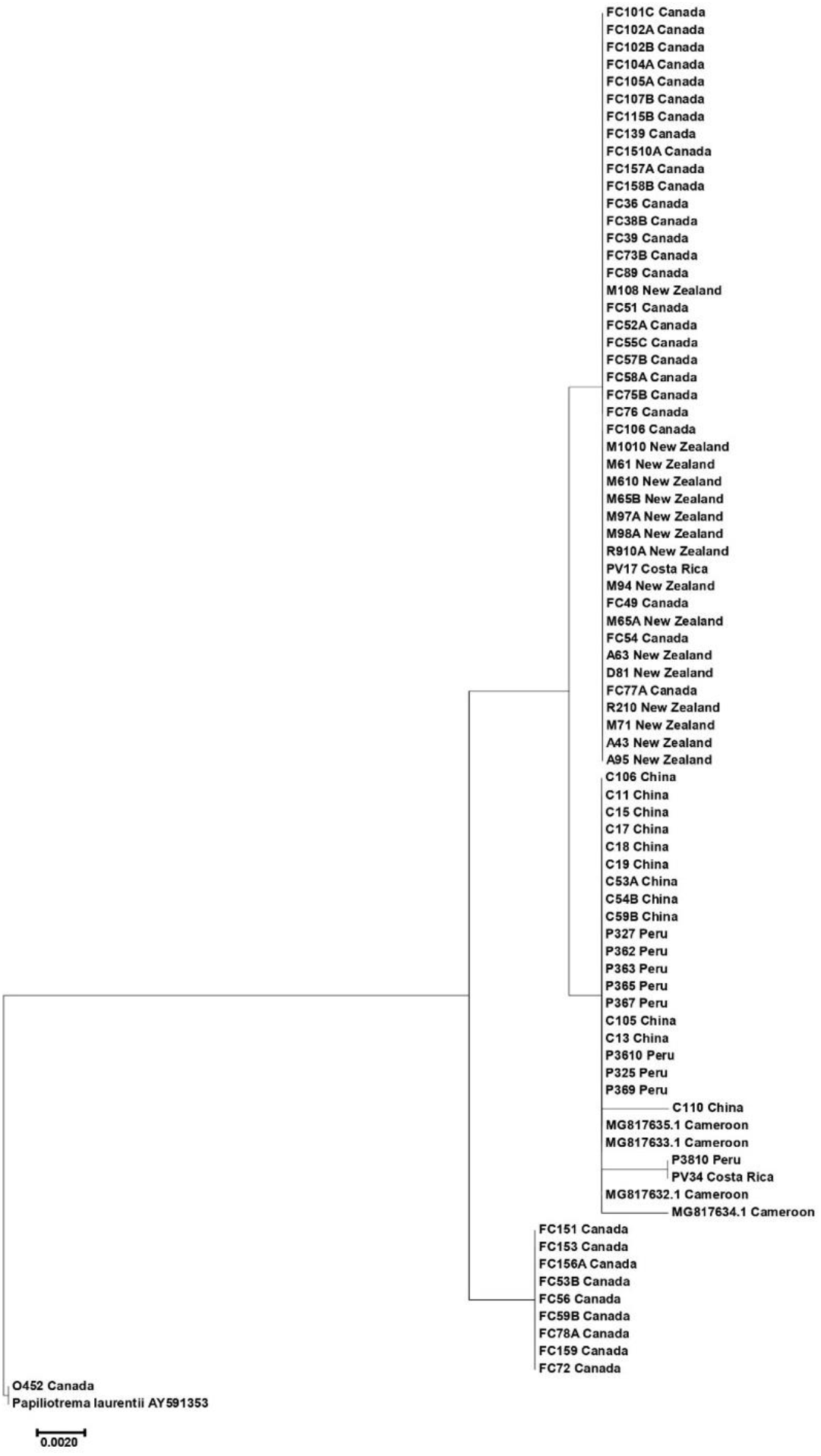
The neighbour-joining tree of *Papiliotreme laurentii* global isolates found in our study. No geographical clustering is observed, suggesting frequent gene flow between populations.

We constructed neighbour-joining (NJ) trees in MEGA7 (Kumar et al., 2016) based on ITS sequences of the four most shared species in our population: *Debaryomyces hansenii* (7 countries), *Papiliotrema laurentii* (6 countries), *Candida tropicalis* (5 countries) *and Torulaspora delbrueckii* (5 countries). The NJ trees highlighted the lack of strict geographical clustering of isolates by country: for an example, most *P. laurentii* isolates found in New Zealand, Costa Rica, China, Peru, and Cameroon had identical ITS sequences and formed a cluster with most Canadian isolates (Figure 7). The NJ trees for the remaining three species can be found in Supplementary Dataset 2. This result is consistent with the hypothesis of recent long-distance dispersals for many of the shared species.

### 3.7 Comparison to culture-independent, metagenomics approach

Our study exclusively used culture-dependent methods to investigate global diversity of culturable soil yeasts. We compared our findings to an excellent study conducted by Tedersoo and colleagues where a metagenomics approach was taken to study global soil fungal diversity (Tedersoo et al., 2014). For the four countries that overlapped between our study and that of Tedersoo et al. (2014), we conducted a detailed side-by-side comparison of results, shown in Table 2. Below we highlight results from two countries.

From Cameroonian soils, Tedersoo et al. obtained 278 fungal OTUs (<u>O</u>perational <u>T</u>axonomic <u>U</u>nits) with species identity determined for 72. Limited information is available for 103 OTUs that were only annotated with a variation of either “fungal_sp” or “uncultured_fungus”. The remaining 103 OTUs were annotated to higher fungal phylogenetic ranks such as Kingdom, Phylum, Class, Order or Family. Our study obtained 110 ITS sequences from Cameroonian soil yeasts. Species identity was determined for 94, genus was established for another nine and the remaining seven were annotated as novel species of the fungal kingdom. Our BLAST analyses revealed no overlapping sequences between the two datasets.

For China, Tedersoo et al. found 572 fungal OTUs, 125 of which were annotated to the species level. The remaining OTUs were annotated to the genus or a higher taxonomic rank. We obtained 230 ITS sequences from Chinese soil yeasts: we established species identity for 209 sequences and genus identity for another three. The remaining 18 sequences, clustered into three groups, were identified as putative novel fungal species. Four sequence overlapped between the two studies and their species annotations were consistent between the two studies.

**Table 3:** Comparison of findings between Tedersoo et al. and current study for four countries (Tedersoo et al., 2014). Tedersoo et al. used metagenomics to determine the diversity of all soil fungi. The current study used culture-dependent methods and fungal DNA barcoding

|  |  | *Tedersoo et al.* |  | *Current study* |  |
| --- | --- | --- | --- | --- | --- |
|  |  | Number | Proportion | Number | Proportion |
| **Cameroon** | Total OTUs found | 272 | 100% | 110 | 100% |
|  | OTUs with species determined | 73 | 27% | 94 | 85% |
|  | Higher taxonomy determined | 52 | 19% | 16 | 15% |
|  | Unannotated | 147 | 54% | 0 | 0% |
| **Canada** | Total OTUs found | 413 | 100% | 261 | 100% |
|  | OTUs with species determined | 104 | 25% | 230 | 88% |
|  | Higher taxonomy determined | 198 | 48% | 31 | 12% |
|  | Unannotated | 111 | 27% | 0 | 0% |
| **China** | Total OTUs found | 572 | 100% | 230 | 100% |
|  | OTUs with species determined | 145 | 25% | 209 | 91% |
|  | Higher taxonomy determined | 235 | 41% | 21 | 9% |
|  | Unannotated | 192 | 34% | 0 | 0% |
| **New Zealand** | Total OTUs found | 269 | 100% | 155 | 100% |
|  | OTUs with species determined | 73 | 27% | 146 | 94% |
|  | Higher taxonomy determined | 87 | 32% | 9 | 6% |
|  | Unannotated | 109 | 41% | 0 | 0% |

## 4. Discussion

Despite being one of the most accessible ecological niches, soil remains an enigmatic source of yeast diversity and ecology. Given that most yeast species are not geographically widely distributed, extensive environmental sampling across diverse regions, habitats and climates is required to uncover new species and diversity patterns. Elucidating global trends and dynamics would also allow us to predict the structure and diversity of soil yeast populations in unsampled locations. Using a set of global soil samples from nine countries in six continents, we address this knowledge gap by characterizing global patterns and predictors of culturable soil yeast diversity. Our study uncovered 134 soil yeast species among 1473 isolates, including 41 previously undescribed species. We identified mean annual precipitation and air traffic volume as significant predictors of soil yeast communities on a global scale. Our findings highlight the influence of both climatic factors and anthropogenic activity on soil yeast populations across the globe.

We found mean annual precipitation to be the strongest predictor of culturable soil yeast diversity across both local and global scales. Previous metagenomic studies have established mean annual precipitation as one of the climatic variables associated with soil fungal diversity (Egidi et al., 2019; Tedersoo et al., 2014). Our results confirm that this trend persists for culturable yeast communities in global soils as well. Vegetation is not a likely mediating factor in the observed positive correlation between precipitation and soil yeast diversity as Tedersoo et al. found plant diversity to be uncoupled from soil fungal richness (Tedersoo et al., 2014). Fungal communities in dry, semi-arid soils contain significantly more Ascomycota fungi than Basidiomycota (Abed et al., 2013; Murgia et al., 2019; Suleiman et al., 2019). We found a reversal of this trend in global soil yeast communities where basidiomycetous yeasts were found to be more prevalent in sampling sites receiving less rainfall (p < 0.05). Some soil-dwelling, basidiomycetous yeasts are known to produce biofilms that allow them to persist in low moisture, oligotrophic conditions (Spencer and Spencer, 1997). Low moisture, and resulting lack of nutrients, could favor cellular structures and metabolic activities of yeasts in one Phylum over the other, creating rainfall-associated global diversity patterns observed in our study. Given our findings, we hypothesize that extreme rainfall and drought events brought on by global warming are likely shifting the established landscape of soil yeast communities. This is especially alarming given the significant presence of pathogenic yeasts we detected in the soils. 15% of all yeast isolates found in our study belong to common pathogenic yeast species capable of causing deadly systemic infections. Altered rainfall patterns, and resulting changes in soil microclimates, could cause outgrowths of pathogenic species and lead to emergence of new fungal infections. With soil ecosystems being a primary source of bacterial and fungal infections, any changes and shifts in soil microbiomes could pose a significant threat to human health.

Each of the nine countries investigated in our study was unique in the composition and structure of its culturable soil yeast population. 73% of the yeast species found in our study (98 out of 134) were specific to a single country. The fragmented nature of soil yeast distributions has been noted in previous studies where only a few species were found to be shared between sampling sites, even within the same region or country (Yurkov, 2018). The nine countries included in our study are separated by thousands of kilometers, with the two closest countries being France and Iceland (2235 km). Geographic isolation was most likely a key factor in limiting the spread and exchange of yeasts between populations, at least until recently when anthropogenic activity has strongly improved the connectivity between countries and continents.

Our findings suggest that human activities can contribute to the changing yeast distribution in soil environments across the globe. International travel has increased exponentially in the past few decades with international tourist arrivals increasing from 25 million in 1950 to a record-high 1.4 billion in 2018 (UNWTO, 2018). The global air transportation network has the small-world property where most countries can be reached from each other via a few flight hops (Wandelt and Sun, 2015). While we found unique yeast species in most localities, 36 yeast species (~25% of all species found) were shared between at least two countries. Countries with more flights occurring between them had more yeast species in common, which implicates human travel as a likely facilitator in the spread of endemic yeasts across geographical borders. The lack of geographical clustering of the most shared species in our population supports gene flow between populations in different countries. The covid-19 pandemic has greatly shifted the political and economical landscape of our planet. Tourism both within and between countries has seen a drastic drop with accompanying tightening of borders between countries. The potential impact of the COVID-19 pandemic on culturable yeast populations remains to be determined.

The current estimate of ~1500 yeast species in existence is almost certainly a significant underestimate (Kurtzman et al., 2011). Both culture-dependent and independent studies routinely isolate novel yeast species from the environment. Our study is one of many recent surveys to find previously undescribed species accounting for as much as 30% of natural yeast populations (Yurkov, 2018), implying that every one of three yeast species recovered from the environment is likely to be a new one. Investigators often turn to natural soils in search of novel yeast strains with commercial and biotechnological potential. A novel strain of *Pichia kudriavzevii* (syn. *Candida kruse*) isolated from soil in a sugarcane filed in Thailand was shown to be more thermotolerant and produce more ethanol than the Thai industrial strain *Saccharomyces cerevisiae* TISTR 5606 (Pongcharoen et al., 2018). Presence of species of the genus *Kazachstania* in mixed cultures of *Saccharomyces cerevisiae* gives rise to fermented wines with diverse aroma profiles: however, *Kazachstania* species are unable to complete fermentation in monocultures (Jood et al., 2017). Discovery of new *Kazachstania* species with more desirable fermentative abilities can aide the full exploitation of this genus in commercial wine fermentation. The thermo and halotolerant yeast *Blastobotrys adeninivorans* is highly useful in a wide range of biotechnological applications including the production of secretory enzymes, as a host for heterologous gene expression and as a biological component in biosensors (Kunze et al., 2017). The metabolic and fermentative capabilities of the novel *Kazachstania* species we found in Peruvian soil and the novel *Blastobotrys* species found in French soil remain to be evaluated.

In recent years, researchers have come to view metagenomics as a valuable tool in the investigation of microbial diversity in complex ecological systems. While high throughput sequencing is crucial in unearthing large-scale patterns at higher taxonomic levels, it fails to be adequately informative on targeted groups of organisms such as yeasts. Our findings indicate that culture-dependent methods can succeed where metagenomics may fail in the study of environmental yeast diversity. Limited information on yeast diversity could be extracted from previous metagenomics studies on global soil fungal diversity (Egidi et al., 2019; Tedersoo et al., 2014). Yeasts do not form a monophyletic group that can be easily identified based on sequences alone. Extracting ITS sequences of known yeasts from large metagenomics datasets is a time-consuming task that requires personnel with advanced knowledge of yeast taxonomy. For potentially novel species, the metagenomic approach would completely fail to identify them as yeasts. The metagenomics approach is largely limited by the current state of knowledge on species taxonomy and annotation. In Tedersoo et al.’s study, species identity was only established for ~25% of the fungal OTUs found in Cameroon, Canada, China and New Zealand: over 30% of the OTUs remained unidentified. Our global collection of soil yeast isolates with identity established and manually validated via ITS sequencing provides a much-needed reference set for future investigators on yeast diversity and taxonomy.

Fungi isolated via culture-dependent methods can be identified as yeasts by morphology and can be further characterized using genomics, metabolomics, and transcriptomics (Xu 2020). Given the relatively low numbers of yeast cells in soil compared to bacteria, mold and other fungi, their DNA can easily escape detection in metagenomics studies, which could explain the lack of overlapping yeast sequences between our study and that of Tedersoo et al. (2014). Our use of culture-dependent methods ensured that the yeasts we found were alive and more likely to be true soil-dwellers as opposed to dead cells temporarily transferred to soil from elsewhere. Selective enrichment and culturing from soil samples in the lab remains the most effective ways of identifying and studying yeasts.

## 5. Conclusions

Our investigation into global patterns in culturable soil yeast diversity reaffirms soil as an important reservoir of environmental yeast species, both known and yet undiscovered. Precipitation emerges as the main predictor of soil yeast diversity across local and global scales. Ongoing global warming crisis and accompanying changes in rainfall could lead to expansion of pathogenic yeasts that already account for a sizable proportion of soil yeast communities. Our findings implicate international travel as a facilitator in the movement of yeast species across borders, with phylogenetic evidence suggesting long-distance gene flow between yeast populations. More environmental sampling is required to further uncover soil yeast diversity, isolates with commercial and biotechnological value and to monitor species that could pose a threat to human health.

## Acknowledgements

We thank the following people for contributing soil samples: Haoran Jia, Yongjie Zhang, Thomas Harrison, Sebastian Harrison, Arshia Kazerouni, and Greg Korfanty for contributing soil samples for this study. HS is supported by an NSERC CGS-D Scholarship. This research is supported by Natural Sciences and Engineering Research Council (NSERC) of Canada (RGPIN-2020-05732) and by the Global Science Initiative Award of McMaster University to JX.

## Author contributions

Study conceived by HS and JX; soil collections were coordinated by JX; lab work performed by HS, YL, RA, AA, and HY; data analyses performed by HS; first manuscript draft written by HS; final draft edited by JX; all authors have read and approved the final version of the manuscript.

